# Interactions of host defense and hyper-keratinization in psoriasis

**DOI:** 10.1101/2021.11.26.469424

**Authors:** Jingwen Deng, Emmerik Leijten, Michel Olde Nordkamp, Sarita Hartgring, Weiyang Tao, Juliette Pouw, Deepak Balak, Rianne Rijken, Runyue Huang, Timothy Radstake, Chuanjian Lu, Aridaman Pandit

## Abstract

**Objectives:** To understand the crosstalk between the host and microbiota in psoriatic skin, using a systems biology approach based on transcriptomics and microbiome profiling.

**Methods:** We collected the skin tissue biopsies and swabs in both lesion and non-lesion skin of 13 patients with psoriasis (PsO), 15 patients with psoriatic arthritis (PsA), and healthy skin from 12 patients with ankylosing spondylitis (AS). We performed transcriptome sequencing and metagenomics profiling on the local skin sites to study the similarities and differences in the molecular profiles between the three conditions, and the associations between the host defense and microbiota dynamic.

**Results:** We found that lesion and non-lesional samples were remarkably different in terms of their transcriptome profiles. Functional annotation of differentially expressed genes (DEGs) showed a major enrichment in *neutrophil activation*. By using coexpression gene networks, we identified a gene module that was associated with local psoriasis severity at the site of biopsy. From this module, we extracted a “core” set of genes that were functionally involved in *neutrophil activation, epidermal cell differentiation* and *response to bacteria*. Skin microbiome analysis revealed that the abundance of *Enhydrobacter, Micrococcus* and *Leptotrichia* were significantly correlated with the “*core network*” of genes.

**Conclusions:** We identified a *core network* that regulates inflammation and hyper-keratinization in psoriatic skin, and is associated with local disease severity and microbiome composition.

## INTRODUCTION

Psoriasis is a chronic inflammatory skin disease which is characterized by erythematous plaques covered with silvery scales. Psoriasis affects approximately 125 million people globally and has multiple comorbidities associated with it, including psoriatic arthritis(1,2). Psoriasis is a complex disease. The development and progression of psoriasis has been associated with several genetic, environmental, and immune factors. Abnormal differentiation of keratinocytes and excessive immune cell infiltration in the skin is considered to be the primary etiology of psoriasis(3).

Extensive evidence implicates the crucial role T cells play in psoriasis(4). However, recent investigations demonstrated that patients with psoriasis have innate immunity disorders, which may have a pivotal impact in the pathogenesis of psoriasis(5–7). In these studies, dysfunction of neutrophils, dendritic cells, and other components of the innate immune system have been reported in patients with psoriasis. However, the precise molecular mechanism underlying innate immunity disturbance and hyper-keratinization has remained inconclusive.

Therefore, it is important to gain better insights into the pathogenesis of psoriasis, especially the innate immune responses in the psoriatic skin environment. In this study, we integrated multi-level data from transcriptome and microbiome based on a cohort of patients with psoriasis and psoriatic arthritis. Our goal was to identify the interaction between the host immune responses and microbiota in psoriatic skin.

## MATERIALS AND METHODS

### Study design

This study was conducted at the University Medical Centre Utrecht (UMCU) and performed in compliance with the Helsinki principles. Ethical approval was obtained from the institutional review board before the recruitment of participants. All participants signed written informed consent before participation.

Skin biopsies, skin swabs, and clinical parameters were collected from a cohort of patients with psoriasis (PsO), psoriatic arthritis (PsA) and ankylosing spondylitis (AS) in a prospective observational study (N=40 patients). The recruitment of participants was performed at the outpatient clinic of the Department of Rheumatology and Clinical Immunology.

### Evaluation of psoriasis severity at the site of biopsy

The Psoriasis Area Severity Index (PASI) scoring method was performed to give a “local PASI” score for the site where biopsy/swab of affected skin was performed (cumulative score of 0-12 based on the total sum: (0~4 redness) + (0~4 thickness) + (0~4 scaling)).

### RNA-seq analysis

Skin samples were derived from 4mm punch biopsies were embedded in Tissue-Tek^®^ and directly snap-frozen in liquid nitrogen, until further processing. Sections were cut (20μm) and dissolved and lysed in RLT plus (+BME) buffer. Samples were vortexed and homogenized by pipets before RNA isolation. RNA isolation was performed using a Qiagen Universal Kit, according to manufacturer’s protocol. Sample were then prepared according to standard operating procedures of GenomeScan BV for sequencing. RNA sequencing (polyA enriched) was performed using Illumina NovaSeq6000 sequencing platform (paired-end, 150 bp) resulting in ~6 Gb (±20 million paired end reads) per sample of Illumina-filtered sequence data.

The reads obtained were aligned to human genome (GRCh38 build 99) using STAR aligner (version 020201) and mapped reads were counted using HTSeq-count (version 0.9.0) (9,10). For all the samples >80% reads unambiguously mapped to the human genome and of those >80% aligned to the annotated genes. DESeq2 (version 1.32.0) was used for the differential expression analysis of different comparisons(11). In differential expression analysis, likelihood ratio test (LRT) was applied for multivariable comparisons or pair-wise comparisons.

### Functional annotation

To identify the biological function of the DEGs and module genes, gene ontology (GO) and Reactome pathway analysis and visualization were conducted using the R package clusterProfiler (version 3.11) (12–14). Gene Set Enrichment Analyses (GSEA) was performed to calculate an enrichment score of gene profiles according to ontology gene sets in the Molecular Signatures Database (MSigDB) C5 collection(15,16). Enrichment scores (ES and NES) were calculated in GSEA. A positive ES indicates the enrichment of genes at the top of the ranked molecular signatures, while a negative ES indicates those at the bottom of the ranked list. The NES is helpful to compare the results across different molecular signatures.

### Estimation of immune cell infiltration

To estimate the immune cell infiltration in skin via gene expression profiles, two computational deconvolution tool xCell is applied. 22 functionally defined human immune cell types were profiled via each tool. Gene signature-based enrichment scores of immune cells were calculated in xCell.

### WGCNA

Filtered, log2CPM normalized gene expression data of lesional samples were used as input for a biweight-midcorrelation signed network which constructed by WGCNA package (version 1.70.3) in R(19). This gene co-expression network was constructed with the following parameters: maxBlockSize=20000, soft power threshold=7, minModuleSize=30, mergeCutHeight=0.2. Network visualization was performed with Cytoscape software (version 3.7.0)(20).

### Gene regulator-target network analysis

Gene regulator-target network for the transcription factors was constructed by using Random Forests methods with RegEnrich R package (version 1.3.0)(21). We analyzed the regulator-target relationship between transcription factors provided by RegEnrich and the DEGs obtained from the psoriatic lesion and non-lesion comparison. RegEnrich provided us ranked list of the regulators along with their significance calculated by integrating the differential expression of regulators, their targets and enrichment analysis(21). In gene regulatory network inferring, both Fisher’s exact test and GSEA were used for the enrichment test.

### *Core network* signature analysis

Gene set variation analysis (GSVA) was used to determine the entirety expression level of *core network*(22). By comparing the GSVA scoring of *core network*, we can estimate the variation in the gene enrichments of the *core network* across different independent cohorts. The GSVA score of *core network* over the psoriatic lesions and controls was measured with R package GSVA (version 3.11). GSVA scores ranged from −1 to 1, with negative scores indicating relative decreases in network expression while positive scores indicated elevations.

### PLS-DA and sPLS-DA classification between PsO and PsA

To identify the most discriminant gene subset between PsO and PsA, we applied both PLS-DA and sparse Partial Least Squares-Discriminant Analysis (sPLS-DA) classificatin modelling on lesional samples with R packages mixOmics (version 6.1.1) (23). To define the number of sPLS-DA dimensions and components, we calculated the overall misclassification error rate and Balanced Error Rate (BER) for each PLS-DA dimension and component selection with maximum distance, centroid distance and Mahalanobis distance. The performance of PLS-DA and sPLS-DA was evaluated by five-fold cross-validation with 100 literations.

### 16S rRNA gene sequencing

Skin microbiota samples were collected by rubbing sterile swabs submerged in SCF-1 buffer for 1 minute on lesional or non-lesional psoriatic skin sites. After swabbing the swab tips were stored in cryovials and stored at −80°C.

Skin swab DNA extraction was performed using the QIAamp UCP Pathogen mini kit automated on the QIAcube. The skin swab was transferred to a tube filled with 0.65ml ATL buffer and incubated for 10 min, 56 °C with continuous shaking at 600rpm. Subsequently 40μl Proteinase K was added and bead beating was performed using the SpeedMill PLUS for 45s, at 50Hz. Samples were then processed according to the manufacturer’s protocol. Approximate amounts of DNA were 1-10 μg/sample. Variable regions V1 and V2 of the 16S rRNA gene were amplified using the primer pair 27F-338R in a dual-barcoding approach (24). DNA was diluted 1:10 prior PCR, and 3μl of this dilution was used for amplification. PCR-products were verified using electrophoresis in agarose gel. PCR products are normalized using the SequalPrep Normalization Plate Kit (Thermo Fischer) and pooled equimolarly.

16S Sequencing of the skin swabs was done using Illumina MiSeq v3 (2×300bp) platform. Demultiplexing after sequencing is based on 0 mismatches in the barcode sequences. The primer sequences used are 27F = AGAGTTTGATCCTGGCTCAG and 338R = TGCTGCCTCCCGTAGGAGT.

### Skin microbiota data analysis

Considering the 16S rRNA data may fail to provide sufficient resolution and accuracy to adequately perform species-level analysis, in our study, only genus-level taxonomy was used for the downstream analysis. To preserve statistical power, only OTUs presented more than 5 reads in at least 20% samples were retained. A total of 140 OTUs in genus level was kept for downstream analysis after the prefiltering. Abundances of microbiota were normalized using the centered log-ratio (CLR) transformation. Principal co-ordinates analysis (PCoA) based on the Jensen–Shannon divergence was performed to visualize differences in the bacterial community structure across samples. Shannon diversity index of microbiota was calculated by vegan packages (version 2.5-5). Microbial abundance was compared across different phenotypes with permutational multivariate analysis of variance (PERMANOVA). Multivariable association between microbiota and phenotypes was analyzed by MaAsLin2 (version 3.13)(25). The circus plot for microbiota-gene association was made by circos (http://circos.ca/).

### Statistical analysis

For clinical characteristics, the chi-square test was applied for non-continuous variables, and analyses of variance (ANOVAs) was applied for the continuous variables. For multiple hypothesis testing, P values were adjusted with the Benjamini-Hochberg method and adjusted FDR P values < 0.05 were the threshold for significant difference. Spearman correlation was calculated between gene expression and microbial abundance. All these statistical analyses were performed using R (version 4.0.3) (http://cran.r-project.org/).

## RESULTS

### Cohort description

The cohort included 13 PsO patients with a dermatologist-confirmed diagnosis of psoriasis in whom concomitant PsA was clinically excluded by a rheumatologist; 15 patients with PsA fulfilled ClASsification of Psoriatic ARthritis (CASPAR) criteria(8); 12 patients with a clinical diagnosis of AS were included as a non-psoriatic reference group, none of which had a history of psoriasis.

The PsO, PsA and AS groups were matched for age, gender, BMI, and smoking. The disease duration for psoriasis, PASI and local PASI scores were in the similar level between the PsO and PsA groups. ESR and CRP as two biomarkers commonly used to detect inflammation, showed no difference between diseases. The clinical characteristics of the participants are shown in Table 1.

**Table 1.** Clinical characteristics of the participants.

|  | **PsO**<br>**(n=13)** | **PsA**<br>**(n=15)** | **AS**<br>**(n=12)** | **P value** |
| --- | --- | --- | --- | --- |
| **Sex** |  |  |  |  |
| Female | 7 (53.8%) | 5 (33.3%) | 3 (25.0%) | 0.302 |
| Male | 6 (46.2%) | 10 (66.7%) | 9 (75.0%) |  |
| **Age (years)** | 41.3 (13.2) | 46.3 (9.32) | 44.2 (8.54) | 0.466 |

|  | **PsO**<br>**(n=13)** | **PsA**<br>**(n=15)** | **AS**<br>**(n=12)** | ***P* value** |
| --- | --- | --- | --- | --- |
| **BMI** | 28.2 (7.79) | 26.9 (4.62) | 25.2 (2.66) | 0.728 |
| **Smoking** |  |  |  |  |
| never | 3 (23.1%) | 4 (26.7%) | 5 (41.7%) | 0.509 |
| past | 5 (38.5%) | 7 (46.7%) | 6 (50.0%) |  |
| present | 5 (38.5%) | 4 (26.7%) | 1 (8.3%) |  |
| **PsO**<br>**Duration (months)** | 24.2 (14.6) | 23.1 (16.1) | - | 0.734 |
| **PASI** | 6.83 (5.40) | 5.05 (4.87) | - | 0.249 |
| **Local PASI** | 5.31 (2.56) | 5.54 (2.15) | - | 0.677 |
| **DMARD past** |  |  |  |  |
| Yes | 12 (92.3%) | 8 (53.3%) | 10 (83.3%) | 0.0433* |
| No | 1 (7.7%) | 7 (46.7%) | 2 (16.7%) |  |
| **UVB past** |  |  |  |  |
| Yes | 7 (53.8%) | 3 (20.0%) | 12 (100%) | <0.001* |
| No | 6 (46.2%) | 12 (80.0%) | 0 (0%) |  |
| **Biologic past** |  |  |  |  |
| Yes | 12 (92.3%) | 14 (93.3%) | 12 (100%) | 0.632 |
| No | 1 (7.7%) | 1 (6.7%) | 0 (0%) |  |
| **ESR** | 9.46 (12.9) | 8.27 (9.61) | 7.82 (7.78) | 0.922 |
| **CRP** | 4.97 (7.30) | 5.04 (6.85) | 4.94 (5.22) | 0.955 |

Mean (standard deviation) are presented for continuous values. Frequencies (proportion) are presented for categorical values. DMARD, UVB or biologic history: usage in past 3 months.
\*Significant at *P* value < .05.

### Transcriptome profile significantly altered between psoriatic lesion and non-lesional skin

We first examined the differences between conditions (PsO, PsA, and AS) and type of skin samples (lesional and non-lesional) using principal component analysis (PCA). The PCA found that the lesional samples from PsO overlaid with the lesional samples from PsA, whereas all the non-lesional samples (PsO, PsA and AS) clustered together (figure 1A). Differential expression analysis showed that lesional samples did not differ between conditions (PsO and PsA) (figure 1C). Similarly, we found minimal differences between non-lesional samples based on their conditions (0 DEGs for PsO vs PsA, and 2 DEGs for PsO+PsA vs AS comparisons) (figure 1C).

**Figure 1.**
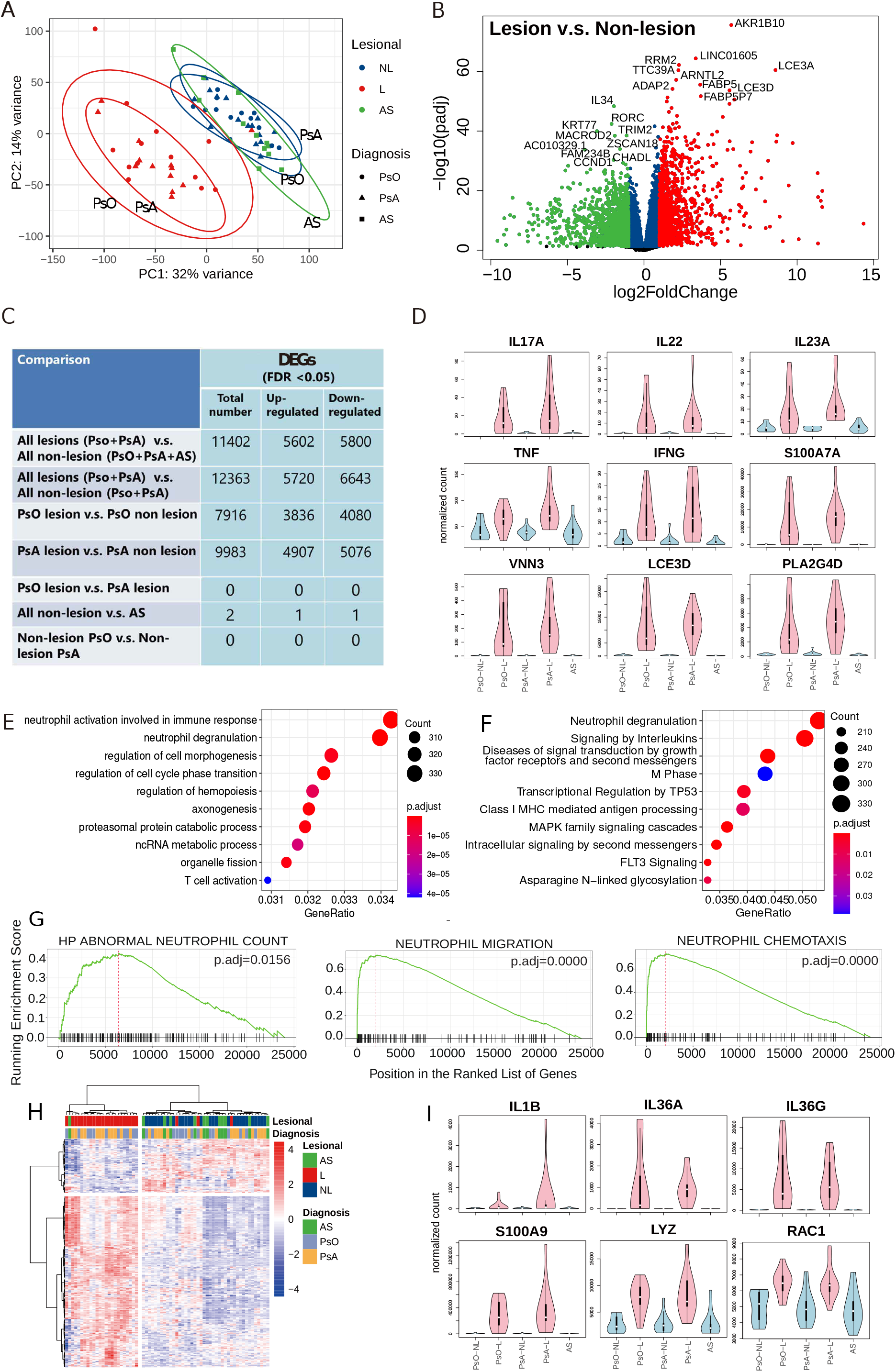
Differential gene expression in psoriatic lesion and non-lesion. **A.** PCA of all samples. **B.** Volcano plot of differentially expressed genes (DEGs) in psoriatic lesion, compared to non-lesion. The red dots are the genes with both false discovery rate (FDR) smaller than 0.05 and log_2_FC greater than 1. The green dots are the genes with both false discovery rate (FDR) smaller than 0.05 and log_2_FC smaller than −1. **C.** Numbers of DEGs in different comparisons. **D.** Expression profile of psoriasis markers DEGs in different cohorts. Gene expression values were normalized with median of ratios method. **E.** Gene ontology (GO) annotation for the DEGs. Top 10 terms are listed. **F.** Reactome annotation for the DEGs. Top 10 terms are listed. **G.** Gene set enrichment analysis (GSEA) of the gene expression profile for the enrichment of neutrophil counts and function in the Molecular Signatures Database (MSigDB) collections C5. **H.** Heatmap of the gene expression profile of neutrophil activation in lesion and non-lesion. **I.** Expression profile of neutrophil markers DEGs in different cohorts.

Since we did not find differences between type of samples from different patient cohorts, we pooled the psoriatic diseases together for the differential expression analysis to further study the etiology of psoriatic lesions compared to non-lesional skin. We found remarkably high number of DEGs (13,700 genes; 6,093 increased and 7,607 decreased) in pairwise analysis of the lesion samples with the non-lesion samples from PsO and PsA (figure 1B, table S1). The commonly used markers of psoriasis(26) were significantly increased/upregulated in lesion samples compared to non-lesional samples (figure 1D). Most of the DEGs we found were consistently increased in lesions of PsO and PsA patient when compared to non-lesional skin samples from PsO, PsA and AS (figure 1C and figure S1A).

To understand the remarkable difference between the transcriptome profiles of lesional and non-lesional samples, we next performed deconvolution analysis using xCell to predict the cell compositions of the different skin samples. In line with previous literature, xCell revealed that different subsets of T cells (bulk CD4+, bulk CD8+, effector memory CD8+ and regulatory T cells) were more abundant in lesional skin compared to non-lesional skin (figure S2). These observations suggested a large immune infiltration in the skin microenvironment.

### Patients with PsA and PsO share a transcriptome profile of skin

Both PCA and DE analysis indicated that the gene expression pattern in lesion of PsO was similar to that of PsA (figure.1A, figure.S3A, tab.1). No gene was identified as DEG between lesional of PsO and PsA with DE analysis (tab.S2). We next asked whether the lesional transcriptome discriminates PsA from PsO in supervised manner. Two supervised learning methods PLS-DA and sPLS-DA were employed to construct classifiers to explore the distinctions between PsA and PsO. PLS-DA is one of the most commonly used methods for supervised classification. It can help to distinguish the differences among samples from different conditions and split the hyperspace of the variables (27). PLS-DA classifier did not provide us a satisfactory of distinction for PsO and PsA (figure. S3B). Considering the noisy variables or non-informative in PLS-DA could affect the interpretability of model and performance of prediction (28). We built another classifier with sPLS-DA method, a sparse approach within the framework of PLS-DA. The performance evaluation of variables in each sPLS-DA dimension showed that top seven variables (LNC-LBCS, AL359962.3, THAP7-AS1, DNAJC8P1, ID2-AS1, GPX1P1 and AL133482.1) in component one combined with top one variable (ZNF396) in component two performed best (figure. S3C). However, most of these eight variables were non-coding RNA (LNC-LBCS, AL359962.3, THAP7-AS1, ID2-AS1, AL133482.1), or pseudogenes (DNAJC8P1, GPX1P1). We applied these eight gene to create the classifier for PsO and PsA. Although sPLS-DA classifier had reduced irrelevant variables and achieve a simpler and more interpretable model (29), it still failed to divide the samples from PsO and PsA completely (figure. S3D, E). Taken together, lesion from patients with PsO and PsA share a transcriptome signature, which supports the concept of a single psoriatic spectrum of disease without major whole-transcriptomic differences. Based on these observations, we combined the lesional samples, non-lesional samples from PsO and PsA as psoriatic lesion group and nonlesion group, respectively. Then we used the DEGs between lesion and non-lesion for the downstream analysis.

### Neutrophil-mediated inflammation in psoriatic lesion

To understand the functional implications of the changes in transcriptomic profiles of lesional skin, we next performed functional annotation analysis (using Gene Ontology (GO), Reactome pathways and GSEA). Using GO annotation, we found that genes attributed to *neutrophil activation involved in immune response* (GO:0002283) and *neutrophil degranulation* (GO:0043312) were significantly enriched in lesional skin (figure 1E). Reactome pathways analysis further supported neutrophil involvement in psoriasis, as neutrophil degranulation (R-HSA-6798695) was the top one pathway enriched in lesional skin (figure 1F). Using the expression profile of 335 DEGs constituting the GO term of *neutrophil activation involved in immune response* clearly separated the lesional samples from the non-lesional and AS samples (figure 1H). Further analysis revealed that more than 75% of these genes associated with neutrophil activation were upregulated in lesional samples.

The GSEA further corroborated that the differences in gene expression profile in lesion skin compared to non-lesional skin were linked to the up-regulation of the cell counts and function of neutrophils. We found significant enrichments of *abnormal neutrophil count, neutrophil migration* and *neutrophil chemotaxis* (figure 1G). We found increased expression of neutrophil associated genes in lesion skin (figure 1I)(30). Using deconvolution analysis, we confirmed that neutrophils were the only myeloid cells that increased in their abundance in psoriatic lesions (figure S2). In summary, multiple analysis indicate that neutrophils are dysregulated in psoriatic lesions.

### Transcriptome alterations were associated with the severity of local lesion rather than the global severity

To obtain clusters of genes (modules) involved in common functions and related to the clinical phenotypes, we constructed the co-expressed network with the gene expression profile of lesion samples using WGCNA. Within this lesional network, we identified eight co-expression modules (black, blue, brown, green, pink, red, turquoise and yellow) (figure 2A). We annotated these eight modules with GO terms and correlated them with clinical parameters (figure 2B, C). We found that disease severity was correlated with the eigengene of five modules (green, yellow, black, pink and red modules; figure 2C). Typically, the severity of lesion where we took the biopsies (local PASI) was more correlated with the gene modules as compared to the global PASI (figure 2D).

**Figure 2.**
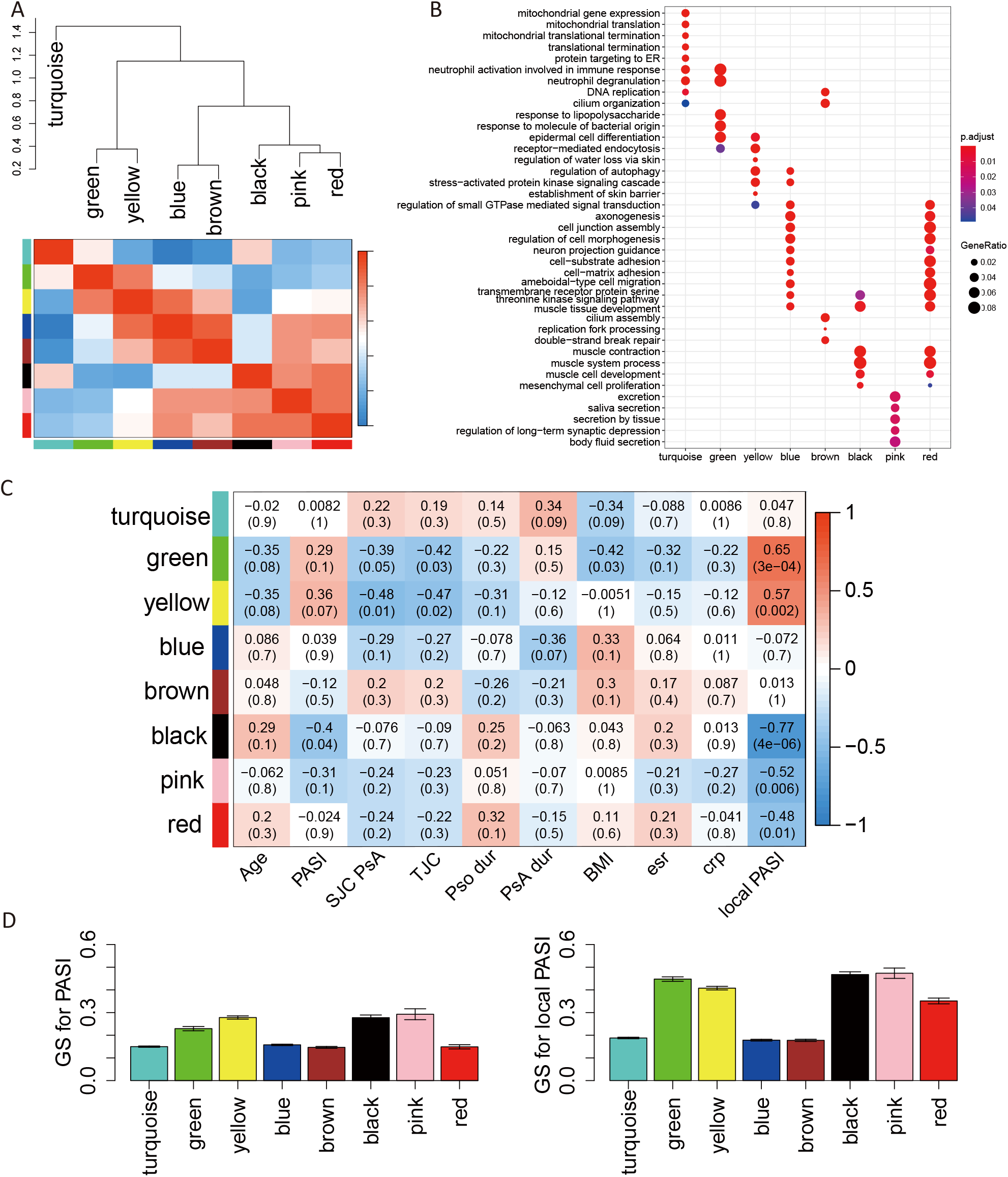
WGCNA construction for gene expression profile and module-trait relationship in lesion samples. **A.** Hierarchical clustering tree (dendrogram) and gene co-expression module definition. in lesion samples. A total eight modules were identified. **B.** Gene ontology (GO) annotation for the modules. **C.** Table of module-trait (clinical parameter) correlations and *p* values. Each cell reports the correlation (and p value) resulting from correlating module eigengenes (rows) to traits (columns). The table is color-coded by correlation according to the color legend. **D.** The significant relationship between modules and phenotypic variables PASI score and local PASI score. GS: gene significance. The results are expressed as the means ± SD.

### Core network was found to correlated with disease severity

We next focused on the local PASI-associated modules (green, yellow, black, pink, red). Green and yellow modules were positively correlated while the other three were negatively correlated with the local PASI (figure 2C). So, we further investigated each gene in green and yellow modules by correlating their expression to local PASI (called gene significance) and using gene network criteria of module membership (a measure to assess if a gene is central or hub of the module and hence important from gene network perspective) (figure S4A, B). The hub genes in green module had higher positive correlations and hence were coupled to the local PASI better than those in yellow module. Moreover, green module was enriched in genes associated with psoriasis-related functional pathways such as *epidermal cell differentiation, the neutrophils activation, neutrophils degranulation, receptor-mediated endocytosis*, and *response to LPS and bacteria* (figure S4C); whereas, yellow module genes were enriched in skin barrier related functional pathways.

We then integrated the knowledge about functionality of the genes from literature and importance of the genes as derived from our data-driven networks. To do so, we extracted the genes that were hub genes in the green gene module and were functionally annotated in psoriasis related GO terms (*epidermal cell differentiation, neutrophils activation and degranulation*, and *response to LPS and bacteria*). By measuring the adjacencies as the edge weights, these hub genes were functionally related and were highly correlated with each other forming a *“core network”* of genes (figure 3A).

**Figure 3.**
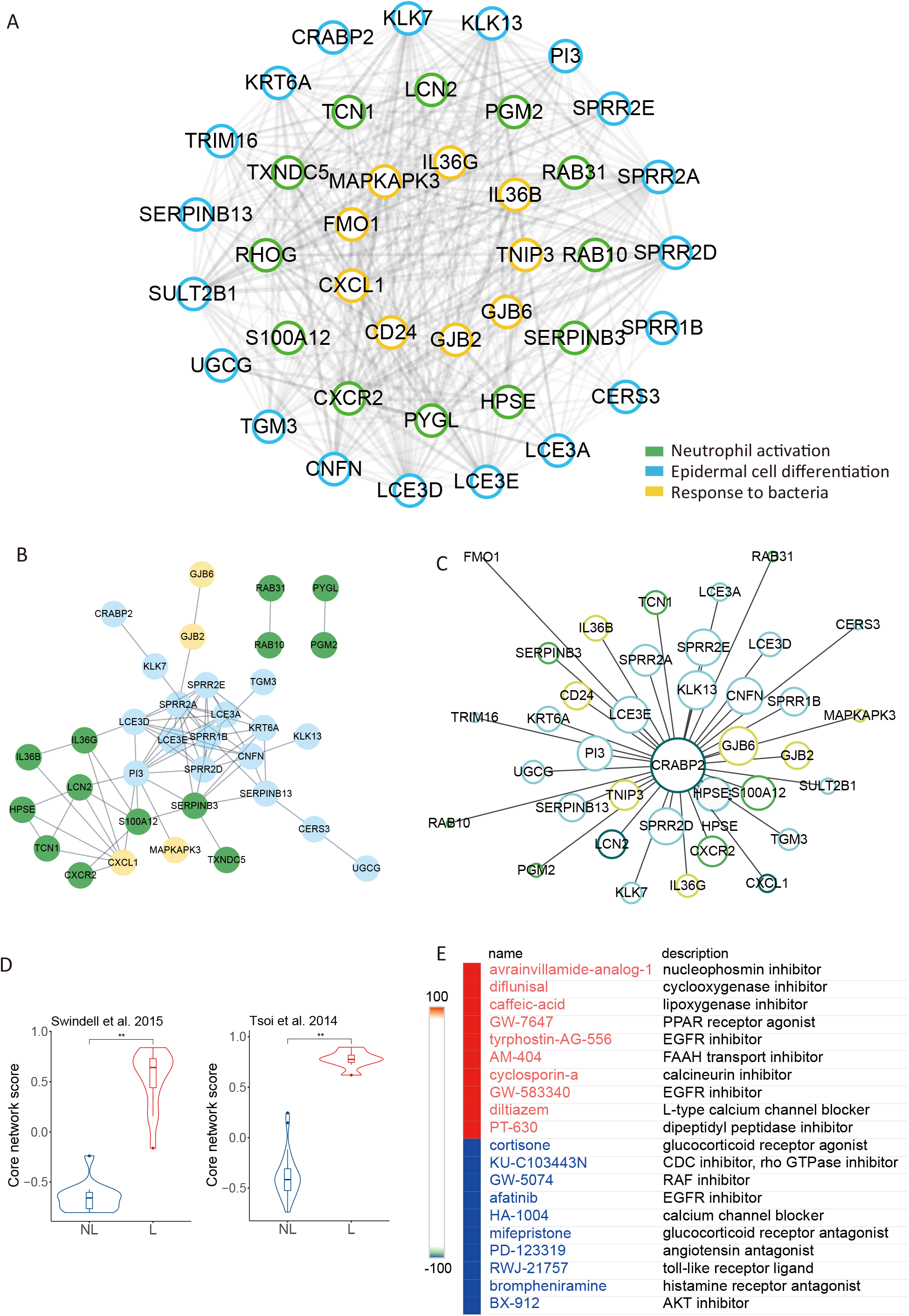
Core network exploration in green module. **A.** Core network in green module: the clusters with the highest adjacency in heatmap (two triangles) were extracted and visualized with cytoscape. Genes for neutrophil activation (light blue), epidermal cell differentiation (green) and response to bacteria (yellow). **B.** PPIs network for core network genes. **C.** Gene regulatory network of CRABP2 and its target genes, only showed the genes overlap with core network. Gene regulatory analysis was based on random forest algorithm. **D.** Core network scores in public datasets (Swindell 2015 and Tsoi 2014). \**p* < 0.05. ** *p* < 0.01. **E.** Drug targets towards the genes in core network. The color was for the median tau score of drugs. The positive score indicated the potential suppress effect of drugs on psoriasis. The negative score indicated the drugs may aggravate the development of psoriasis.

We considered that the genes in the *core network* could be highly correlated with each other as: a) they are physically interacting with each other, or b) they are regulated by the same set of transcriptional regulators. So, we first explored if the genes in the *core network* are known to have physical interactions using the STRING database(31) and found that 75% of the *core network* genes were involved in physical protein–protein interactions (figure 3B). We further used RegEnrich to perform the gene regulatory network analysis. A large proportion (36/40) of genes in *core network* were overlapped with the target genes of CRABP2 in gene regulatory network, which implied that the genes in *core network* not only represented co-expression pattern but also the regulator-target relationship (figure 3C). Interestingly, both protein-protein network analysis and gene regulatory network analysis revealed that *core network* genes CRABP2, a member of a family of specific carrier proteins for Vitamin A, as an important regulator of genes within the *core network* and of genes differential in lesional samples (figure 3B, 3C, S5A). All these evidences indicated that the genes in *core network* had more interactions among themselves than what would be expected for a random set of genes with co-expression pattern, and that these genes were at least partially biologically connected to each other as a network.

To ascertain the reproducibility of our *core network*, we used two independent cohorts of psoriatic skin samples from publicly available datasets. We used GSVA method to estimate the strength of *core network* signature in a sample. In this method, a higher score means higher expression of *core network* genes. The *core network* scores of lesional skins were consistently higher than those of the non-lesional group across two independent datasets (figure 3D). The expression profiles of these two independent datasets showed that the *core network* genes were upregulated in lesional samples, with sufficient intensities (figure S5B).

To discover the well-annotated drug–gene interactions relevant to medical decision-making based on our *core network*, we further investigated the potential drugs targeted to the *core network* in CLUE database (32). The results demonstrated that avrainvillamide (nucleophosmin inhibitor), diflunisal (cyclooxygenase inhibitor), caffeic-acid (NF-kB pathway inhibitor), Cyclosporin-a (calcineurin inhibitor) among others were targeting the *core network*, implying their potential to be utilized for psoriasis treatments. Interestingly, Cyclosporin-a is an established treatment for psoriasis was also defined as a suppression to *core network* (figure 3E).

### Skin microbiota influenced disease severity and associated with the *core network*

As the functional annotation of green module indicated the response to bacteria was involved, we integrated the skin microbiome with the skin transcriptome in our analysis. Similar to the transcriptome profile, PCoA plot showed that the samples were clustered based on skin type (lesional or non-lesional) and not on clinical phenotype condition (figure 4A). PsO showed an increase in diversity, compared to AS. Lesional pools were more diverse than non-lesional pools and AS (figure 4B). The diversity of microbiota in skin was positively correlated with psoriasis duration and age (figure 4C). To figure out the distinction between lesion and non-lesion, we additionally carried out a non-parametric multivariate analysis of variance of clinical phenotypes on the psoriatic skin microbiota. Our results indicated that multiple clinical factors (age, sex, use of DMARD, etc.) influenced the differential abundance analysis (table S3). After correction for these confounding effects, we got three genera differentially enriched between PsO and PsA, five genera significantly associated with PASI and four with local PASI (figure 4D).

**Figure 4.**
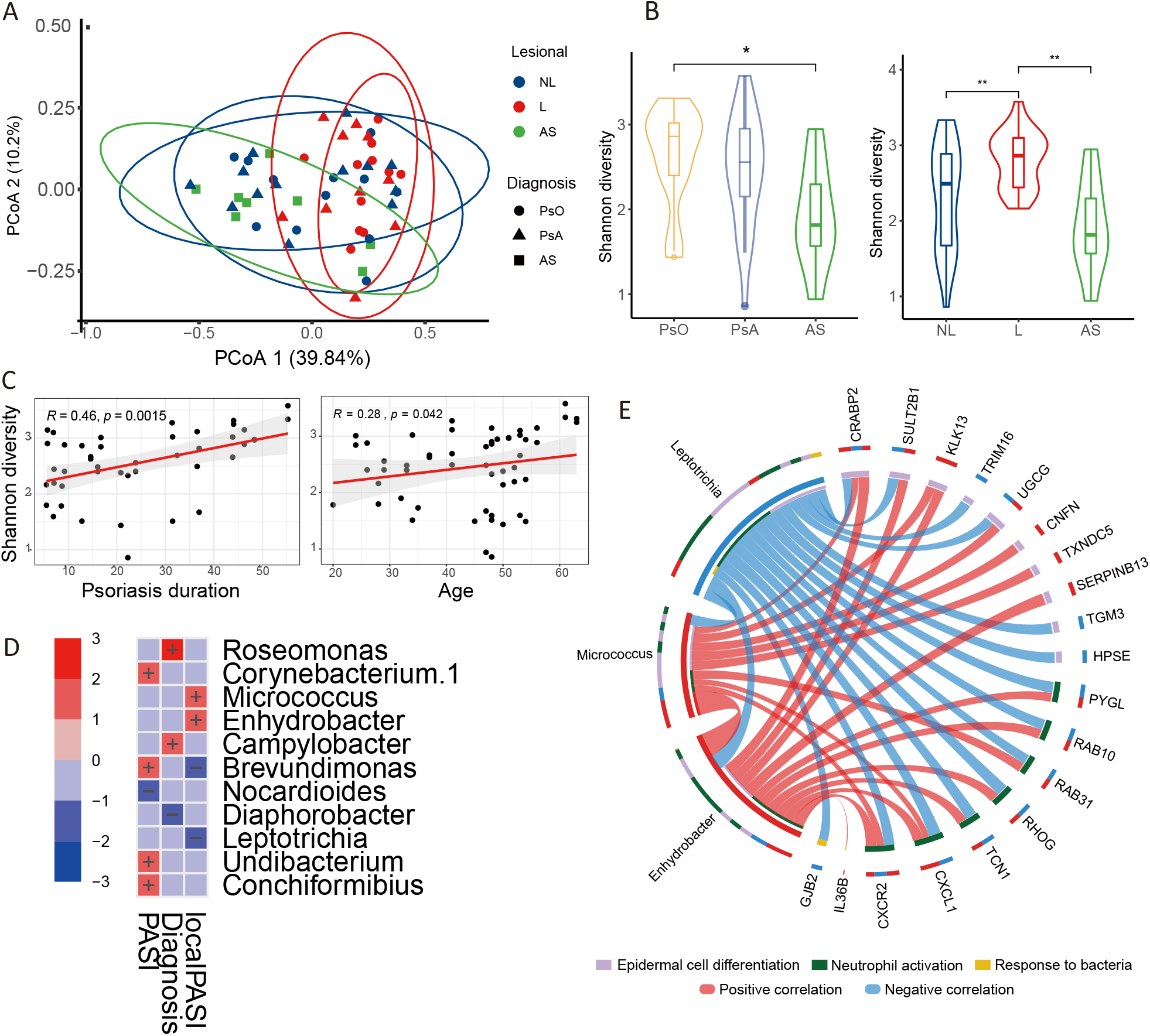
Microbiota profile and host-commensal interaction. **A.** Principal coordinates analysis (PCoA) of skin microbiota based on OTUs in genus level. **B.** Shannon diversity of skin microbiota in different cohorts. **C.** The association between the Shannon diversity and psoriasis duration (left) or age (right). **D.** Heatmap of the significant associations between skin microbiota abundance and clinical parameters for all samples (upper) and for only lesional samples (lower). Color for association Significance (*p* < 0.05). Sign “+” and “−” for coefficient. **E.** The association between the abundance of *Enhydrobacter, Micrococcus, Leptotrichia* and core network genes. Size of ribbon size encodes *R* values. All ribbons showed were for correlations with statistical significance. * *p* < 0.05. ** *p* < 0.01.

We subsequently evaluated whether the four local-PASI-associated genera were also related to the core gene network. Interestingly, the abundance of *Enhydrobacter, Micrococcus* and *Leptotrichia* were strongly correlated with the expression of genes in *core network* (figure 4E). The gene expression of CRABP2, CXCL1, CXCR2 was associated with the abundance of all these three genera. Moreover, *Enhydrobacter, Micrococcus* and *Leptotrichia* were also significantly correlated with each other (figure 4E), implied the potential microbe-microbe interactions in skin.

Overall, host gene-microbiome interactions in psoriatic skin were revealed by integrating the profile of disease severity associated genera and gene expression of *core network*.

## DISCUSSION

The precise pathogenesis of psoriasis remains incompletely understood. Recent studies indicate that host-environment interactions in the microenvironment of skin may play a central role in the development of psoriasis. Here, we found that neutrophil dysregulation was a major factor contributing to the distinct transcriptomic profiles of psoriatic skin lesions. We identified a *core network* of genes associated with disease severity, inflammation and hyper-keratinization in psoriasis. These *core network* genes were associated with the abundance of skin microbiota, highlighting the interactions between local environment and immunopathogenesis of the skin.

Figure 5 summarizes our major findings. We identified *core network* of genes that were dysregulated in psoriatic lesions (figure 3 and 5). Several of these genes have previously been associated with psoriasis such as SERPINB3, LCN2, LCE3A/D/E and/or associated with interaction with local microbiota such as CXCL1, CXCR2, PI3, SERPINB13, IL36, LCN2. For example, SERPINB3 has been shown to produce Pso p27, an immunogenic protein complex that is produced via post-translational modifications specifically in psoriasis (39). LCN2 is a known clinical marker for itch in psoriasis (40). LCE3A/D/E can be induced when the skin barrier is disrupted and are strongly expressed in psoriatic lesions (38).

**Figure 5.**
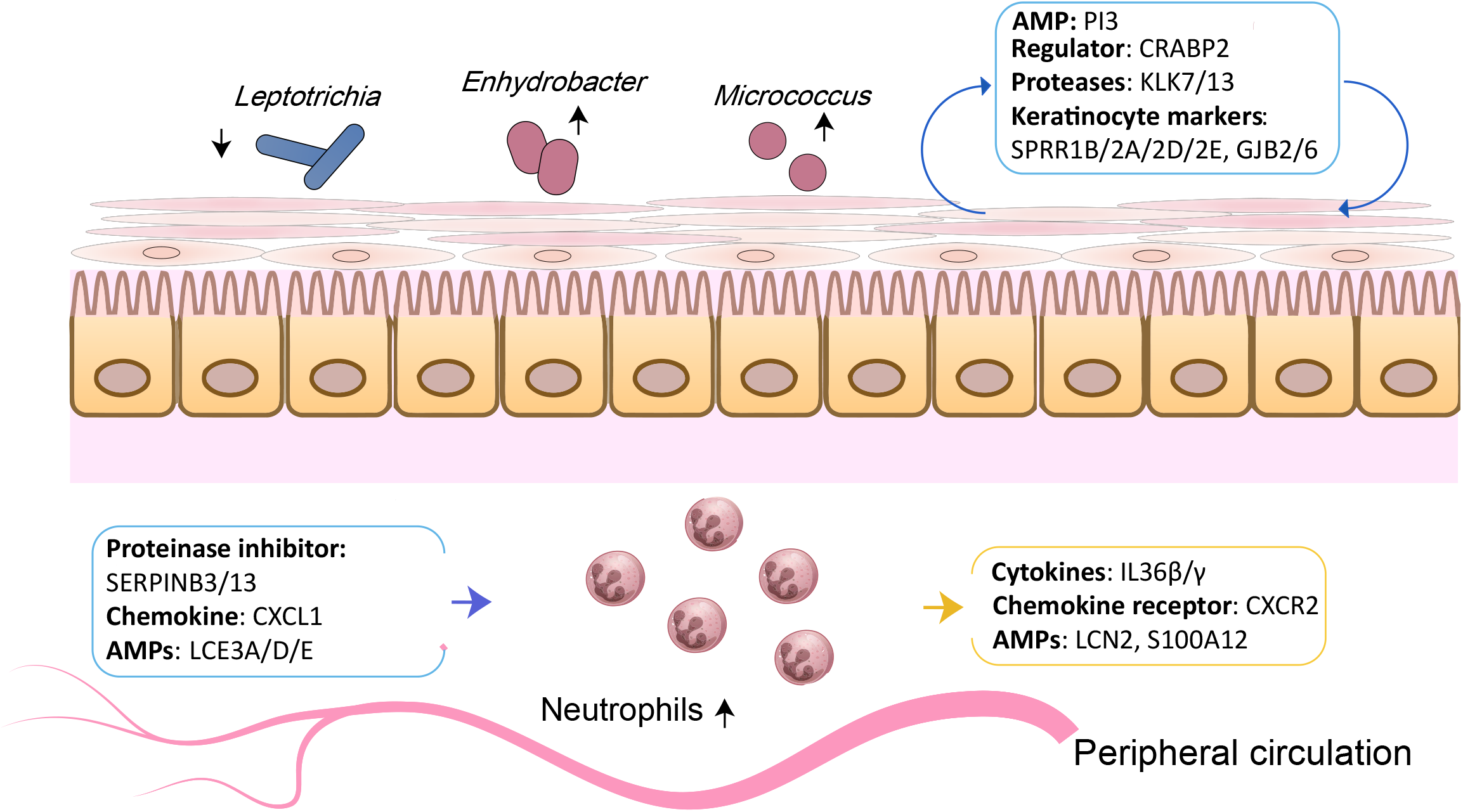
Interactions of host defense and hyper-keratinization in psoriatic lesion.

Interestingly, many of genes in the *core network* were strongly correlated with the alterations in the local microbiota (*Enhydrobacter, Micrococcus*, and *Leptotrichia*) indicating that microbial imbalances in skin environment is associated with specific changes in the transcriptomic profiles (figure 4E and 5). These microbial imbalances induce the defensin-like antimicrobial activity of keratinocytes including the secretion of SERPINB3/13 and antimicrobial peptides (PI3 and LCE3A/D/E). SERPINB3/13 are reported as alternative activators of IL-1α and IL-1β which have very important roles for antimicrobial host defense (41). The defensin-like antimicrobial activity of keratinocytes has been shown to recruit neutrophils from circulation to skin. For example, overexpression and secretion of CXCL1 by keratinocytes facilitates skin homing of neutrophils by binding to CXCR2 receptor. Activated neutrophils then produce proinflammatory mediators such as cytokines (IL36β/γ) (42) and antimicrobial peptides (LCN2, S100A12) in the skin. IL36β/γ enhance IL-1α level and amplifying the defensin-like antimicrobial activity(43), ultimately induced the expression of SERPINB3 (44). LCN2 binds and sequesters the iron-scavenging siderophores to attenuate bacterial growth(45).

These proinflammatory mediators stimulate the keratinocytes leading to hyper-keratinization ultimately leading to overexpression of keratinocyte differentiation markers (SPRR1B/2A/2D/2E, GJB2/6) (46–48), regulators (CRABP2) (49) and proteases (KLK7/13) (50). Interestingly, PI3 was also secreted by keratinocytes to hinder the neutrophil chemotaxis in skin.

In summary, we identify specific host-microbe interactions and core set of dysregulated genes that drive the pathogenesis of psoriasis. We also identified potential target genes and relevant drugs molecules that can be repurposed to psoriasis but would need further exploration for clinical translation. Further research is needed to determine if modulating the skin microbiota composition and/or therapeutically targeting the *core network* can restore skin homeostasis in psoriasis. One of the major strengths of our study is the availability of multiple types of data from the same individual. However, our study has limitations, which includes the relatively low number of samples. As part of the study protocol, we included skin samples from AS patient and assumed these biopsies to be healthy as these patients did not have clinical skin disease at the site of biopsy.

## ACKNOWLEDGMENTS

We would like to thank the patients for participating in the study. We would like to thank the clinical study team (Nienke Kleinrensink, Nanette Vincken, Anne Karien Marijnissen, Anneloes van Loo, Karin Schrijvers, and Joke Nijdeken).

## AUTHOR CONTRIBUTIONS

Conceptualization: AP, JD, EL

Methodology: AP, CL, JD, EL

Data curation: JD, WT, RR

Formal analysis: JD, EL

Funding acquisition: TR, EL, AP, SH, JD

Project administration: EL, SH, AP, RH

Supervision: AP, CL, TR

Resources: MON, JP, DB, EL, AP

Writing – original draft: JD, EL, AP

Writing – review & editing: AP, JD, EL, CL, MON, SH, WT, JP, DB, RR, RH, TR

## DATA AND MATERIALS AVAILABILITY

The skin transcriptome data used in the analysis is available in GEO database (GSE186063).

