## Supplemental figure 1 for "Interactions of host defense and hyper-keratinization in psoriasis"

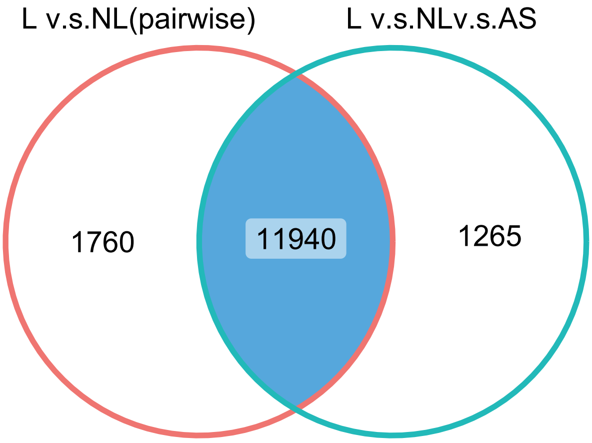


**Figure S1.** Venn diagram showed the overlap of DEGs across the L v.s. NL pairwise comparison and L v.s. NL v.s AS comparison.
