## Supplemental figure 2 for "Interactions of host defense and hyper-keratinization in psoriasis"

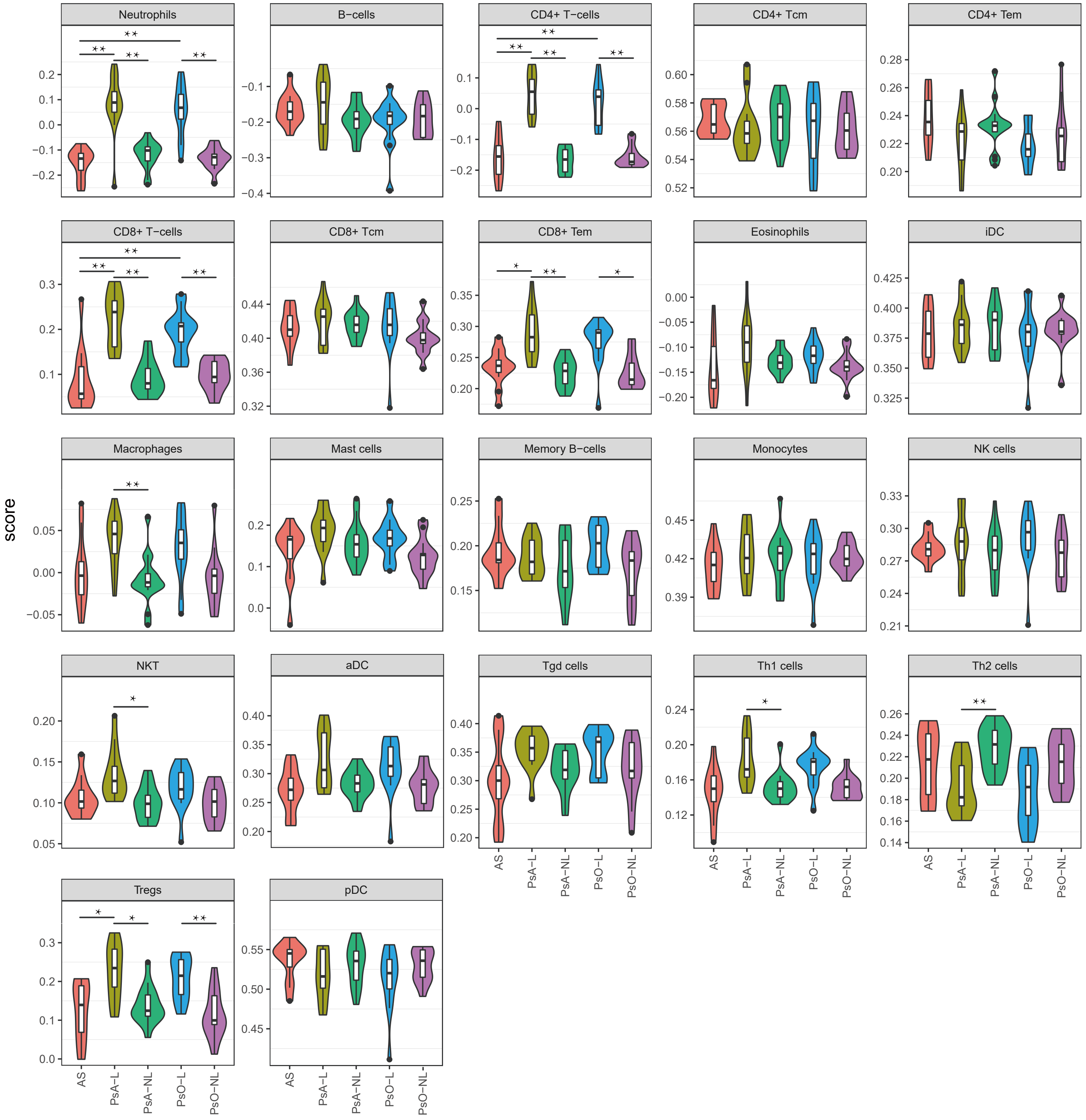


**Figure S2.** Infiltration score of immune cells in skin predicted by deconvoluting bulk gene expression profile with xCell .
