## Supplemental figure 3 for "Interactions of host defense and hyper-keratinization in psoriasis"

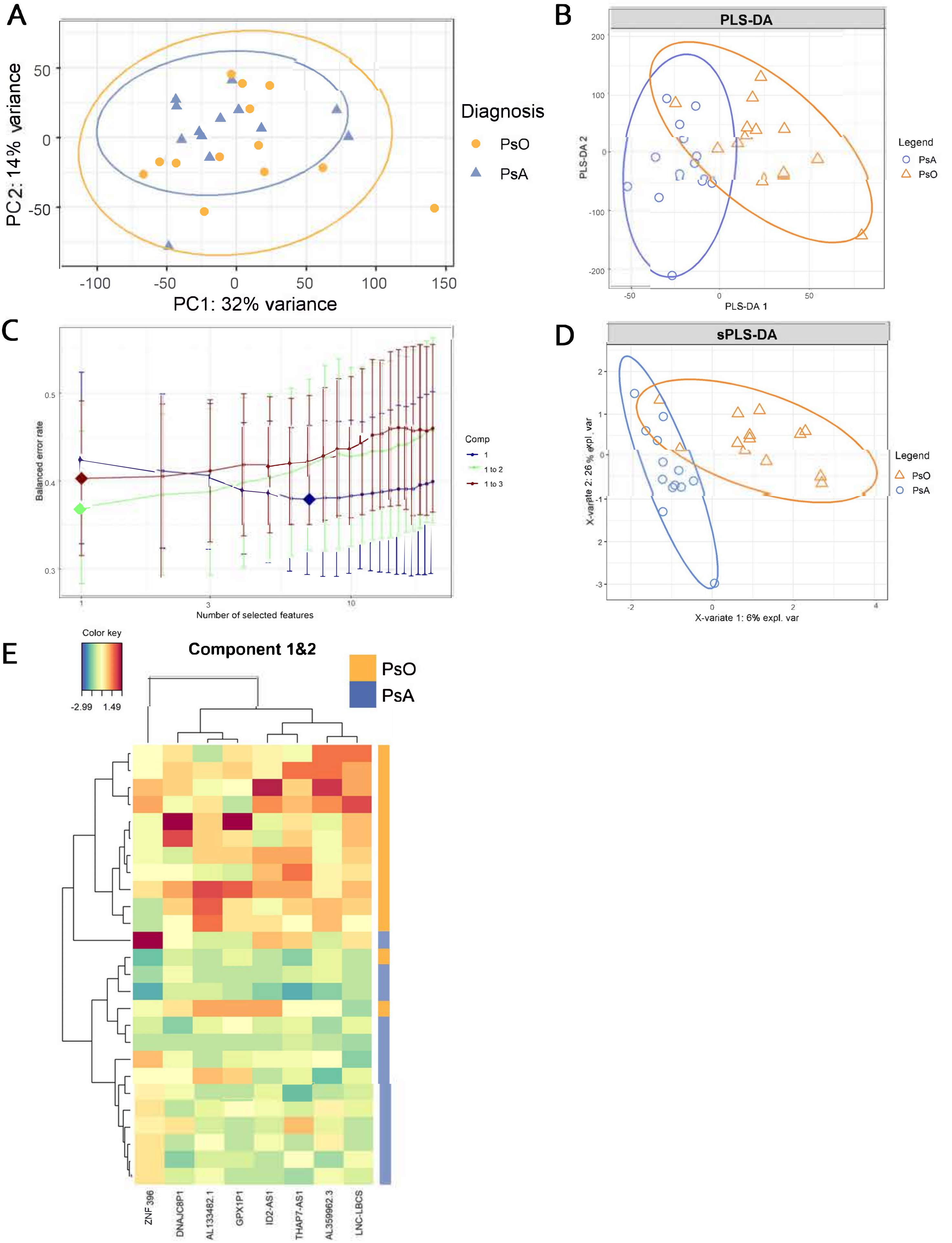


**Figure S3**. Exploration of the deferentially expression profile between lesion samples of PsO and PsA.
**A.** PCA of lesional samples.
**B.** Supervised analysis with PLS-DA.
**C.** Performance of feature selection with sPLS-DA method.
D. Supervised analysis with sPLS-DA.
E. Clustered heatmap of lesional samples in rows with selected features in columns.
