## Supplemental figure 4 for "Interactions of host defense and hyper-keratinization in psoriasis"

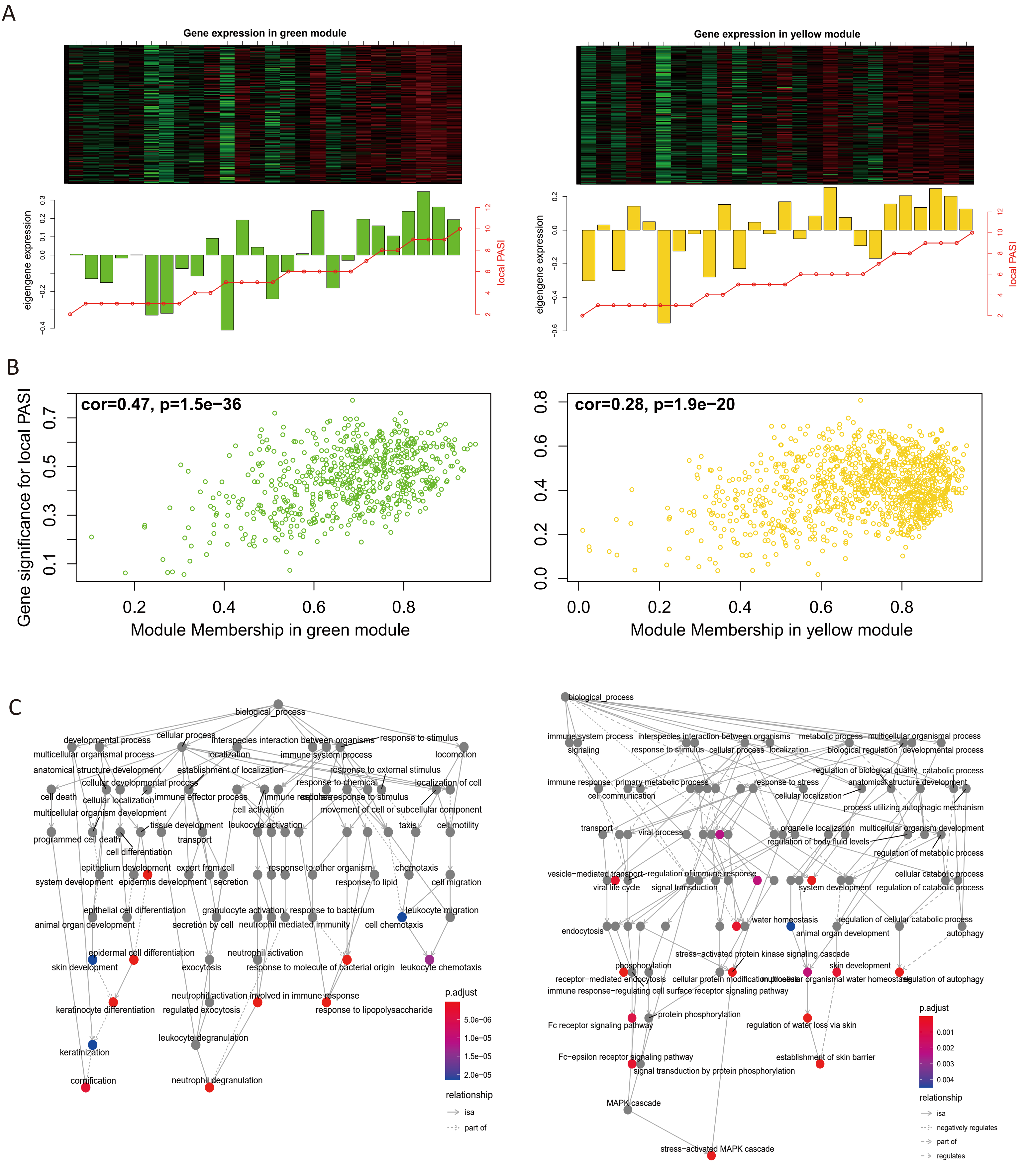


**Figure S4.The patterns of green module and yellow module.**

**(A)** The correlation between local PASI and green module (left), or yellow module (right). The upper panel is the heatmap plot of the expression of module genes (rows) across the samples (columns) which are ranked by local PASI from low to high. The vertical bands are the module eigengene value of samples. The red line is for the local PASI scores of corresponded patients.

**(B)** Visualization of gene significance (GS) for local PASI vs. module membership (MM) in green (left) and yellow (right) modules.

**(C)** The GO network of module genes, for green module (upper), and yellow module (lower).
