## Supplemental figure 5 for "Interactions of host defense and hyper-keratinization in psoriasis"

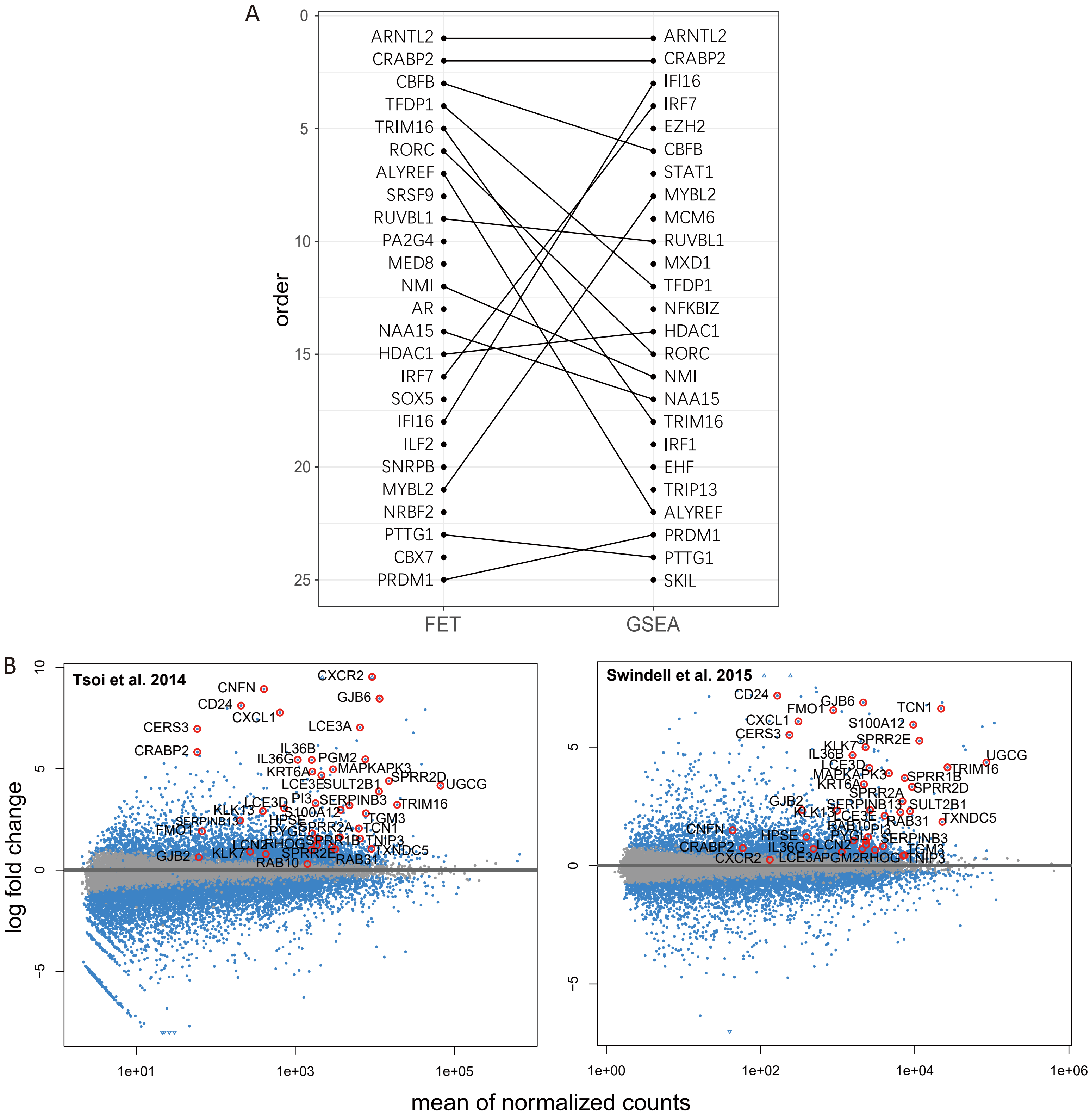


**Figure S5. Validation for core network.**

**(A)** The plot of regulator orders. The regulators identified in gene regulatory network inferring were ordered by their corresponding importance score measured by FET and GSEA methods. The Same regulators are linked by lines.

**(B)** MA plots of two independent datasets. The blue dots were DEGs. The core network genes were labelled with red circles and their names.
